# Identifying mutant-specific multi-drug combinations using Comparative Network Reconstruction

**DOI:** 10.1101/2020.12.17.423240

**Authors:** Evert Bosdriesz, João M. Fernandes Neto, Anja Sieber, René Bernards, Nils Blüthgen, Lodewyk F.A. Wessels

## Abstract

Inhibition of aberrant signaling with target inhibitors is an important treatment strategy in cancer, but unfortunately responses are often short-lived. Multi-drug combinations have the potential to mitigate this, but to avoid toxicity such combinations must be selective and the dosage of the individual drugs should be as low as possible. Since the search space of multi-drug combinations is enormous, an efficient approach to identify the most promising drug combinations and dosages is needed.

Here, we present a pipeline to prioritize promising multi-drug combinations. We performed a limited set of drug perturbations in an isogenic cell line pair with and without an activating PI3K mutation, and recorded their signaling states and cell viability. We used these data to reconstruct mutant specific signaling networks and map the short term signaling response to longer term changes in cell viability. The resulting models then allowed us to predict the effect of unseen multi-drug combinations, at arbitrary drug-concentrations, on cell viability. Our initial aim was to find combinations that selectively reduce the viability of the PI3K mutant cells, but our models indicated that such combinations do not exist for this cell line pair. However, we were able to validate 25 of the 30 low-dose multi-drug combinations that we predicted to be anti-selective. Our pipeline thus enables a powerful strategy to rapidly map the efficacy and possible selectivity of drug combinations, hence significantly speeding up the pace at which we can explore the vast space of combination therapies.

## Introduction

The dependency of tumors on activated signaling pathways results in therapeutic responses to inhibitors that block pathway activity [1]. However, resistance to such targeted inhibitors inevitably develops [2, 3]. Combinations of two targeted inhibitors can give more lasting clinical benefit, but resistance nonetheless emerges [4, 5]. Combining more than two drugs might further extend the duration of the response [6], but toxicity becomes a major concern when multiple drugs are combined at their maximum tolerated dose. Recently, we found that partial inhibition of three or four kinases by combining Multiple drugs at Low Dose (MLD) is surprisingly effective in receptor tyrosine kinase (RTK) driven tumors in multiple cancer types [7]. It prevents the development of resistance, and it is well tolerated by mice. Others have also shown the potential of multi-drug (low dose) combinations in pre-clinical [8–11] and clinical [12, 13] settings.

These findings warrant further exploration of multiple-drug combination strategies. This will require a systematic way to explore promising drug combination treatments, including optimizing the dosing of the different drugs. The combinatorial explosion of the search space — there are more than 2 million possible 4-way combinations of the 89 (as of 2020 [14]) FDA approved targeted inhibitors, and 24 billion if each drug is to be tested at 10 different concentrations — means that in-vitro testing of all combinations is infeasible. Computational approaches are required to prioritize promising combinations.

Recently, Nowak-Sliwinska and collaborators presented a “Feedback Systems Control” approach to explore the search-space of possible multi-drug combinations [10, 15, 16]. While this approach is promising, the method does not optimize for selectivity and the obtained results lack a mechanistic underpinning, making it hard to assess to what extent the results will generalize. Another promising approach is building mathematical models of cellular signaling, based on a limited set of perturbation experiments [17–24]. However, current approaches suffer from two major shortcomings. First, only a very limited number of such modeling approaches focus on the difference between cells with different mutation profiles [17, 25], which is critical for optimizing selectivity. Second, how inhibition of oncogenic signaling affects cell viability, and specifically to what extend short-term signaling response is informative for longer-term cell fate, remains underexplored [23, 24, 26].

We therefore set out to establish and validate a combined experimental and computational pipeline to prioritize multi-drug combinations and their dosing based on mathematical models of drug response (Figure 1). Importantly, we aimed to find combinations that are selective for cells with a particular oncogenic driver mutation. To isolate the effect of the mutation, we used an isogenic cell line pair with and without a mutation. Specifically, we used MCF10A, a cell line derived from epithelial breast tissue [27], and an isogenic clone with the activating PI3K^H1047R^-mutation knocked in under its endogenous promoter [28]. We measured the response of the MAPK and AKT pathway and cell viability after drug perturbations, and used the measurements to build mutant specific signaling networks models using Comparative Network Reconstruction, a method we recently developed [17]. In addition, we found that non-linear model combing the response of phospho-ERK and phospho-AKT is highly predictive for cell viability, despite the fact that signaling response and cell viability are measured on completely different time-scales of hours and days, respectively. Combining the model of mutant signaling with the cell viability model allowed us to simulate the effect of any multi-drug combination at any concentration and thus to prioritize promising combinations. Our models indicated that no drug combination would likely be selective for the PI3K-mutant cells. To nonetheless validate our computational approach, we proceeded to predict which low-dose, multi-drug combinations were likely to be anti-selective, i.e. reduce the viability of the parental cells more strongly than that of the PI3K-mutants. Experimental validation showed that 25 of the 30 combinations that we predicted to be anti-selective indeed had a significantly stronger effect in the parental cell than in the PI3K-mutant cells.

**Figure 1:**
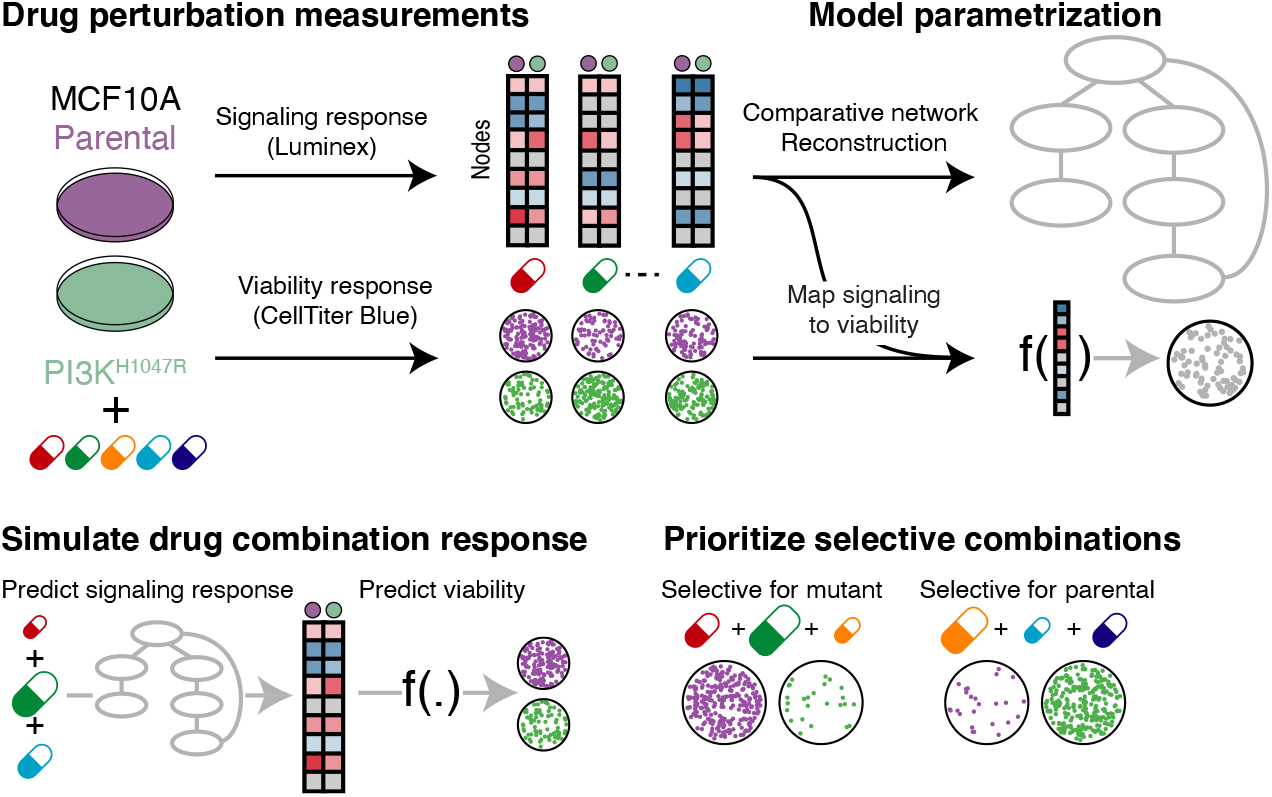
Overview of pipeline to prioritize promising selective low-dose multi-drug combinations. **Top:** MCF10A parental and PI3K^H1047R^ cells are treated with inhibitors targeting the MAPK and AKT pathways. The signaling and cell viability responses are measured and used to build mutant specific models of signal-transduction networks and to parametrize the relationship between signaling response and cell viability. **Bottom:** These models are used to simulate the response to unobserved multi-drug combinations, at arbitrary concentrations, of the signaling networks and how this affects cell viability. In this way, low-dose multi-drug combinations that are likely selective for a particular cell type can be prioritized.

## Results

### The signaling and viability response to drug perturbations in MCA10A parental and PI3K^H1047R^ mutated cells

To test how oncogenic mutations affect signal transduction networks and their downstream effects on cellular phenotypes such as cell viability, we used the MCF10A cell line [27] and an isogenic clone with the activating PI3K^H1047R^ mutation knocked in under its endogenous promoter [28]. As expected, in the PI3K^H1047R^ cells the baseline signaling activity of AKT and PRAS40, both downstream of PI3K, is elevated, but the other signaling nodes do not show significant differences in activity (Figure 2A). In the absence of drug perturbations, PI3K^H1047R^-mutant MCF10A cells have a comparable growth rate as their parental cells [28]. Dose response curves of selected PI3K and the MAPK pathway inhibitors showed subtle differences in sensitivities between the parental and the mutant cells (Figure S1A).

**Figure 2:**
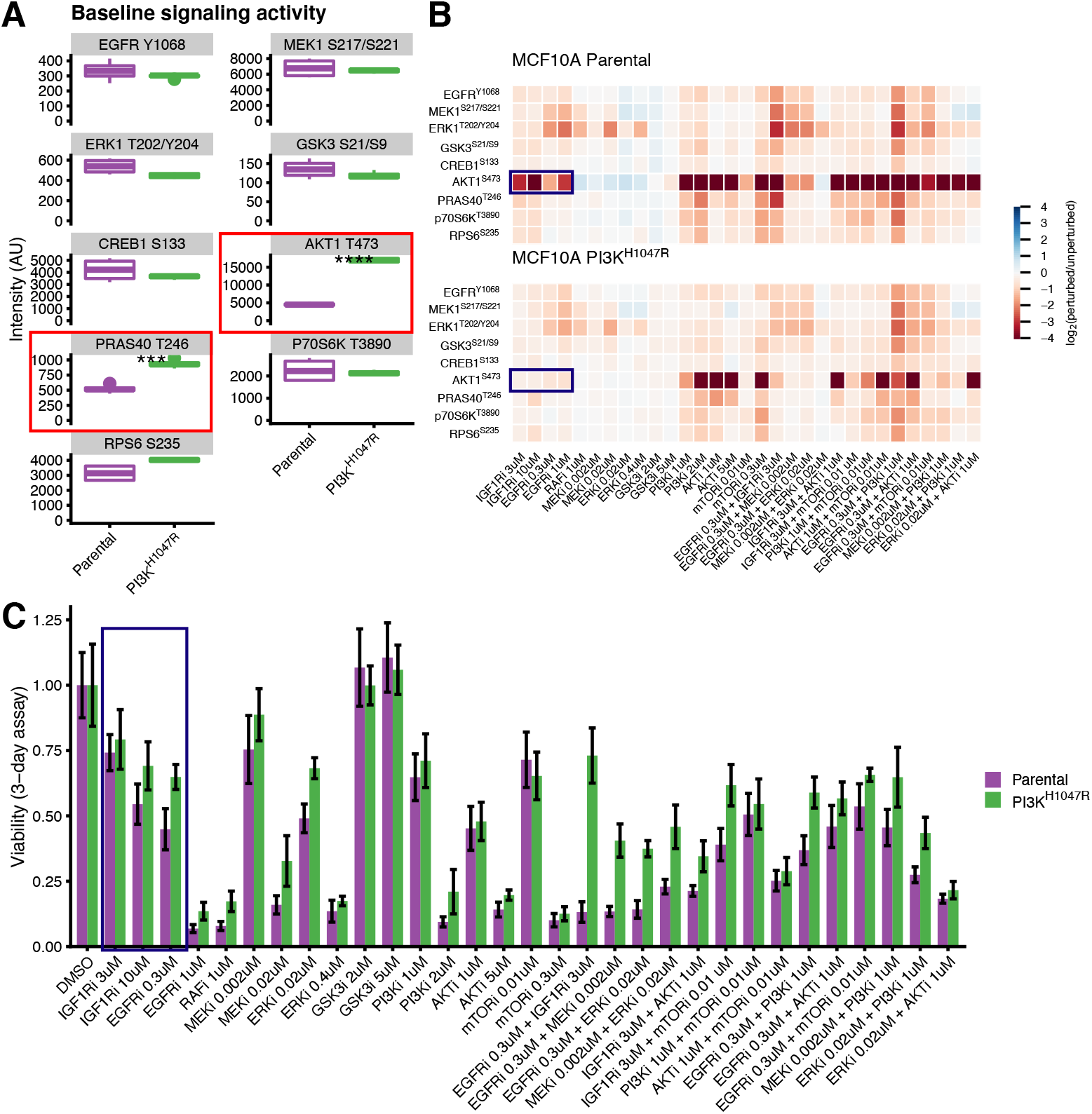
Profiling signaling and viability response of MCF10A parental en PI3K^H1047R^ cells to drug perturbations. **A.** Node activity in the unperturbed cells. Most nodes have similar activity in the parental and PI3K^H1047R^ cells, except AKT and PRAS40 (highlighted) which are downstream of PI3K. **B.** Heatmap representing log_2_-fold changes of the signaling nodes upon drug perturbation compared to DMSO controls. The response of the parental and PI3K^H1047R^ cells are highly correlated, with some exceptions such that of AKT1 upon growth-factor receptor inhibition (highlighted). Signaling response is measured after 2 hours of drug treatment. The color scale is capped between −4 and 4 for visualization purposes. **C.** Cell viability under the same drug treatments as reported in panel B. Both cell lines show a similar response profile. The strong differences in AKT response to growth-factor receptor inhibition translate into mild differences in cell viability (highlighted). Cell viability is measured after 3 days of drug treatment. Error-bars represent standard deviations.

To study how the signaling of these cells respond to drug-perturbations and if the PI3K^H1047R^ mutation influences this, we perturbed both cell lines with inhibitors of the PI3K and MAPK pathways, and selected 2-drug combinations of these. Single drugs were tested at two different concentrations, corresponding roughly to their IC_50_ and IC_90_ values (except RAFi, which was only tested at IC_90_) and drug-combinations were tested with both drugs at their IC_50_ values, to obtain a total of 34 different perturbations. We measured the response after two hours of drug treatment (log_2_ fold change relative to DMSO control) of nine main nodes in the PI3K and MAPK signaling pathway using a multiplexed luminex assay to obtain more than 600 signaling drug-response measurements (Figure 2B, Table S1 and S2). We selected the two hour time-point because this is the timescale for phospho-AKT (pAKT) to reach quasi steady state after PI3K-pathway inhibition [29] (Figure S1B). Luminex quantification showed excellent concordance with Western Blots (Figure S1C). In addition we measured the effect on cell viability using CellTiter Blue (Figure 2C, Table S3 and S4). Generally, the differences in both signaling response and cell viability between the parental and PI3K-mutant cells were subtle but consistent. For instance, while the responses of the signaling nodes of the parental and PI3K^H1047R^ cells are strongly correlated (Figure 2B and Figure S1D), pAKT shows a strong negative response to growth-factor receptor inhibition (EGFRi or IGF1Ri) that is nearly absent in the PI3K^H1047R^ mutant cells (Figure 2B, highlighted). However, this results in only mild differences in cell viability between the cell lines (Figure 2C, highlighted).

### Network reconstructions identify relevant differences between parental and PI3K-mutant cells

To establish how the PI3K^H1047R^ mutation affects the signal transduction network, we used the drug-response measurements to perform Comparative Network Reconstruction (CNR) [17] of the MAPK and AKT pathways of both cell lines. CNR is a method that we have recently developed to reconstruct and quantify signaling networks and identify the most important quantitative differences between two or more cell lines. Prior knowledge about the network topology can be included, but the algorithm can also propose edges to be added to the network. The edge-weights are interpreted as the percent change in the downstream node activity in response to a 1% change in activity of the upstream node. Importantly, by penalizing differences between cell line models, CNR identifies which edges are quantitatively different between the two cell lines.

We used the canonical MAPK and PI3K pathway interactions as prior information, and added 4 edges that were proposed by the CNR algorithm based on hyperparameter selected in a leave-one-out cross validation loop (Figs 3A and S2A). The targets of some inhibitors were not measured in our panel. These were modelled as affecting the first downstream target that was measured. For instance, since both IGF1R and PI3K are not measured in our panel, IGF1R inhibition was modelled as targeting AKT1 directly. The model gave a good fit to the data (Pearson correlation = 0.91) (Figure 3B). To assess the significance of this fit, we compared the residuals of the model to 1000 models with the same number of randomly selected edges. Each of these 1000 random models had a worse fit than our model (*p* < 0.001, Figure 3C).

**Figure 3:**
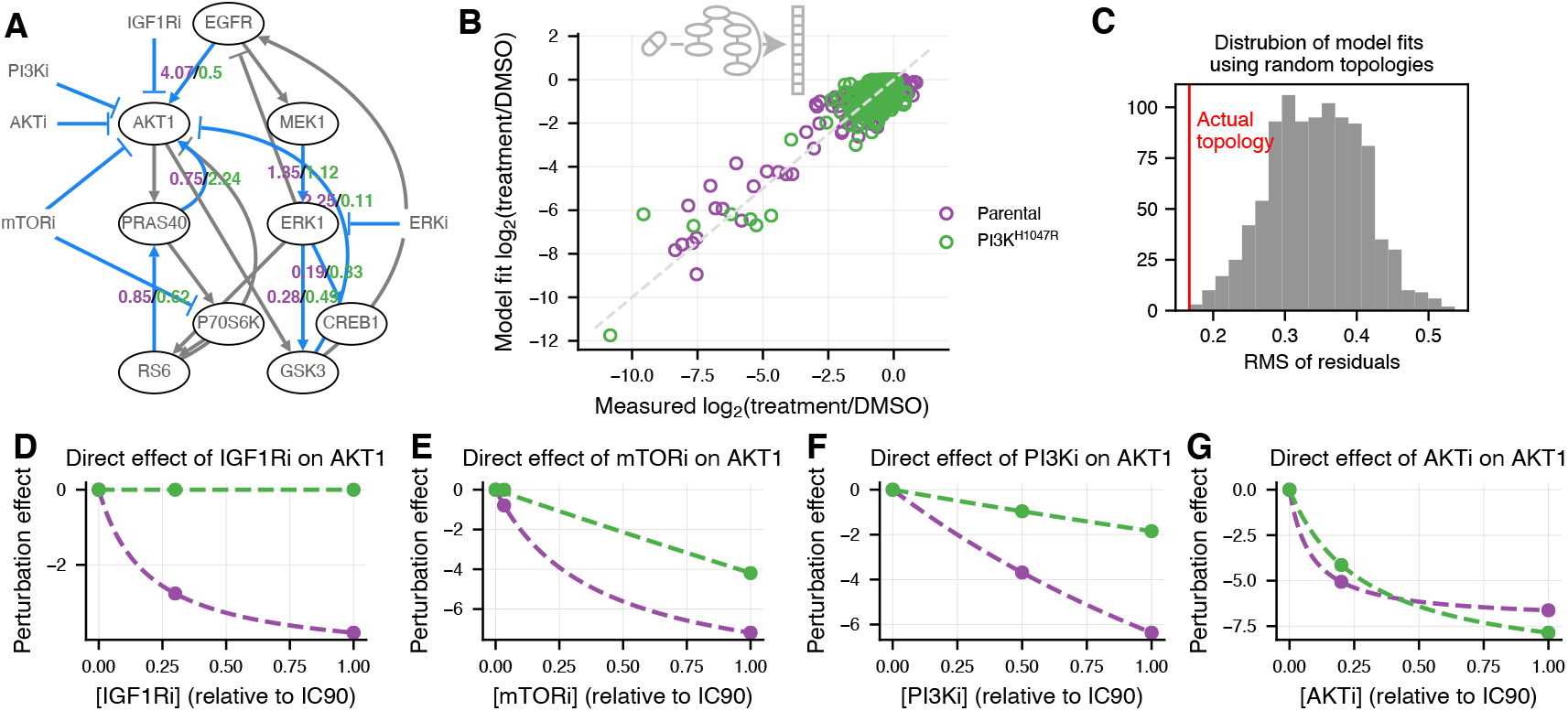
Mutant specific network reconstructions show expected differences. **A.** Comparative Network Reconstruction (CNR) of MCF10A parental and PI3K^H1047R^ cells. Edges and direct perturbation effects that differ between the two cell lines are highlighted in blue. Gray edges do not differ between the cell lines. Edge strengths of the differing edges are represented by the purple (parental) and green (PI3K^H1047R^) numbers. Ovals indicate nodes. For visualization purposes, only direct perturbations effects that differ between the cell are indicated. As expected, the most and the strongest differences between the cell lines are located close to AKT in the network (note that PI3K is not measured). **B.** Comparison of network model fit with measured signaling response shows that the network model can explain the signaling response data well (Pearson correlation 0.91). **C.** Distribution of the root mean square of residuals of models optimized using a random topology (gray), compared to that of the actual model used (red). All 1000 random topology models had the same number of edges as the actual model, and for all 1000 the fit was worse than for the actual model. **D-G.** The estimated direct effect of IGF1R (C), mTOR (D), PI3K (E) and AKT (F) inhibition on AKT activity as a function of applied inhibitor concentration. Points indicate the estimated effects of the concentrations used in the CNR reconstruction, the dashed lines indicate the interpolated curves between these points (c.f. Materials and Methods, Equation 5). IGF1R, PI3K, and mTOR inhibition were modelled as directly affecting AKT because their actual targets were not measured.

CNR aims to identify the most relevant differences between cell lines by penalizing quantitative differences. These differences can be either the edges in the network, or the strength of inhibition of a drug to its direct target. This way, we identified 13 relevant differences between the parental and PI3K^H1047R^ cells. These differences are highlighted in blue in Figure 3A. The numbers next to the edges indicate the edge-strengths of the parental (purple) and PI3K^H1047R^ (green) cells. (For visualization purposes, only the strength of edges that differ between the cell lines are indicated. For full model visualization, see Figure S2B). The differences in target-inhibition strength between the cell lines are shown in Figure 3D-G and Figure S2D. We assessed the significance of the identified differences by comparing the residuals of our model to that of 1000 models with the same number of randomly selected differences. None of the random models had a better fit to the data (Figure S2C), indicating that the identified differences are, indeed, the most relevant differences.

As expected, most of the identified differences are located close to AKT in the network (Figure 3A, note that PI3K is not measured). Specifically, in the PI3K^H1047R^ cells, AKT is less sensitive to changes in EGFR and unresponsive to IGF1R inhibition (Figure 3A and D), which is consistent with PI3K being constitutively activated. Additionally, AKT is less responsive to PI3K and mTOR inhibition (Figure 3E,F). At the IC_50_, AKT is also less sensitive to AKT inhibition, but when AKTi is applied at its IC_90_, the PI3K^H1047R^ cells show a larger response (Figure 3G). This last observation might be explained by the higher baseline AKT activity of PI3K^H1047R^ cells, since if AKT activity is reduced to a similar absolute level, the fold-change of AKT in the mutant is higher.

In order to predict the signaling-response to drugs combined at arbitrary concentrations, we parameterized the relation between target inhibition and drug concentration using the direct target-inhibition estimates for drug *k* on node *i* for the IC_50_ and IC_90_ we obtained from the network reconstructions (c.f. Materials and Methods, Equation 5). The dashed lines in Figs 3D-G and Figure S2C indicate the curves we parametrized in this way.

### Short-term signaling response is informative for long-term cell viability

To prioritize multi-drug combinations, the short term response of the signaling network to a drug perturbation needs to be related to its longer term effect on cell viability. Important open questions here are: Is the short-term signaling response predictive to longer term cell viability? If so, which signaling outputs are most predictive, and what is their relation? The association between the individual node-responses and cell viability were moderate even for the most strongly associated nodes, phospho-AKT (pAKT) and phospho-ERK (pERK), which had a Pearson correlation with cell viability of 0.36 and 0.42, respectively (Figure 4A). The responses of all other nodes also correlated somewhat with cell viability (Figure S3A), but clearly no single node alone is a good predictor for cell viability.

**Figure 4:**
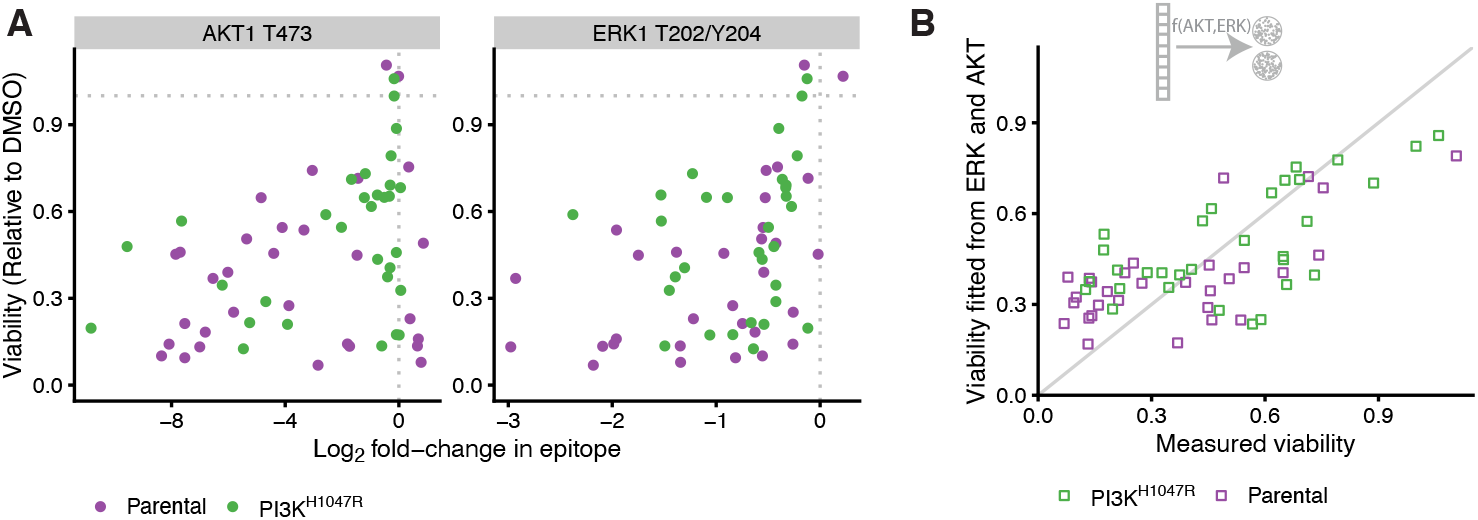
Short term signaling response is predictive for longer term cell viability. **A.** Scatterplot of cell viability against log_2_-fold changes in AKT (left panel) and ERK (right panel) activity in response to drug treatments. The Pearson correlations are 0.36 and 0.42 respectively. **B.** Scatterplot of model fit against measured cell viability based on a model where both ERK and AKT response are used to explain cell viability (c.f. Materials and Methods, Equation 4). The Pearson correlation between fit and measurement is 0.71.

We therefore investigated whether a model combining the response of multiple nodes described the cell viability data better. To this end, we first fitted a linear model using either all nodes or only the response of pERK and pAKT. Both models gave a reasonable fit to the data, but there was a clear structure in the relation between the residuals and the fitted values (Figure S4A and B), indicating that a non-linear model might be more suitable. To test this, we fitted a number of biologically motivated non-linear models relating the combined response of pAKT and pERK to cell viability. These non-linear models that all have the property that cell viability goes to 0 if either pERK or pAKT are fully inhibited (c.f. Materials and Methods, Equation 4). The biological assumption behind this is that both ERK and AKT activation are required for cell survival and growth.

To select the best model, we compared the standardized residuals of the model fits and the L2-norm of the residuals in a leave one-out cross-validation loop of the different models (Table 1). All non-linear models had clearly better performance than the linear models, despite having equal or less free parameters. While overall the predictions of the different non-linear models were fairly similar (with Pearson correlations of their predictions between 0.92 and 0.99, Figure S4D), a Michaelis-Menten like model of the following form

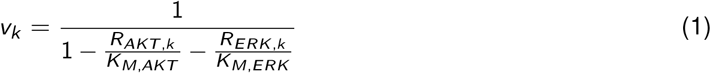

had the overall best performance on both metrics and shows no clear structure in the residuals (Figure S4C). Here, *v_k_* is the cell viability and *R_ERK,k_* and *R_AKT,k_* are the log_2_-fold changes of pERK and pAKT upon drug treatment *k*. The parameters *K_M,ERK_* and *K_M,AKT_* can be interpreted as the log_2_-fold changes of pERK and pAKT that cause 50% inhibition of cell viability. They differ slightly between the two cell lines, but the bootstrapped 95% confidence intervals strongly overlap (Figure S4E), so we do not want to overinterpret these differences. Importantly, this model gave a good fit to the data (Figure 4B), with a Pearson correlation between fitted and measured viability of 0.71.

**Table 1:** Comparison of goodness of fit of functions relating signaling response to cell viability. *σ* is the mean residual standard error of the model fitted to the full data. The L_2_-norm is calculated over predictions made in a leave-one-out cross-validation loop. The non-linear models predict viability *v* from the log_2_-fold change of pERK and pAKT (*R_ERK_* and *R_AKT_*) whereas the linear models fit the inhibition (1 − *v*). The table is ordered from best to worst fit.

| Function | Type | $\sigma$ (Model fit) | L<sub>2</sub>-norm (LOOCV) |
| --- | --- | --- | --- |
| $v_k \sim 1 / \left( 1 - \frac{R_{AKT,k}}{K_{M,AKT}} - \frac{R_{ERK,k}}{K_{M,ERK}} \right)$ | Non-linear | 0.20 | 0.042 |
| $v_k \sim 1 / \left[ \left( 1 - \frac{R_{AKT,k}}{K_{M,AKT}} \right) \cdot \left( 1 - \frac{R_{ERK,k}}{K_{M,ERK}} \right) \right]$ | Non-linear | 0.20 | 0.045 |
| $v_k \sim 3 / \left( 1 + 2^{-\frac{R_{ERK,k}}{K_{M,ERK}}} + 2^{-\frac{R_{AKT,k}}{K_{M,AKT}}} \right)$ | Non-linear | 0.21 | 0.048 |
| $v_k \sim 4 / \left[ \left( 1 + 2^{-\frac{R_{ERK,k}}{K_{M,ERK}}} \right) \cdot \left( 1 + 2^{-\frac{R_{AKT,k}}{K_{M,AKT}}} \right) \right]$ | Non-linear | 0.21 | 0.048 |
| $1 - v_k \sim \sum_{i \in \text{nodes}} \beta_i R_{ik}$ | Linear | 0.21 | 0.061 |
| $1 - v_k \sim \beta_{AKT} \cdot R_{AKT,k} + \beta_{ERK} R_{ERK,k}$ | Linear | 0.28 | 0.093 |

Together, these results indicate that short-term signaling response is informative for longer-term drug response, that pAKT and pERK are the most informative readouts, and that the relation between signaling response and viability is non-linear.

### Prediction and validation of selective multi-drug-combinations

We then combined the network models (Figure 3A) with the parametrization of the signaling-viability model (Equation 4) to simulate the effect on cell viability of unseen 3-drug combination at unseen drug-concentrations. When applying this model to the training data, the Pearson correlation between measured and fitted cell viability was 0.78 (Figure 5A). We used this model to prioritize multi-drug combinations and their dosing that maximize the selectivity, defined as the difference in viability between the parental and the PI3K^H1047R^ cells.

**Figure 5:**
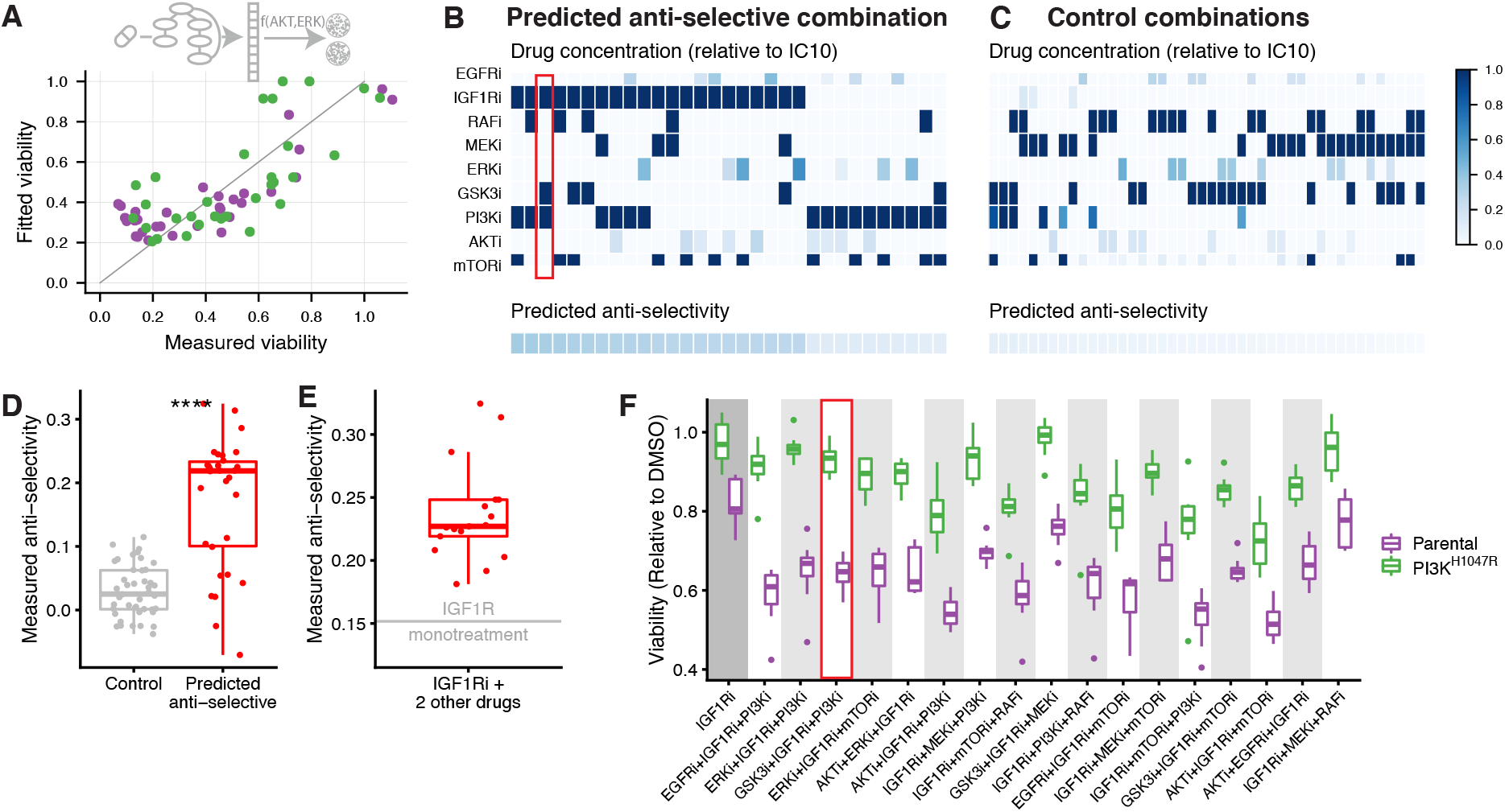
Experimental validation of anti-selective drug combination predictions. **A.** Scatterplot comparing full model fit (network model combined with signaling response-viability mapping) to the training data. The Pearson correlation between fit and measurement is 0.78. **B-C.** Overview of drug combinations that we predicted to be anti-selective (B) and non-selective (C) based on this model. Drug concentrations are color-coded relative to their IC_10_. Bottom row indicates predicted anti-selectivity (defined as the difference in viability between PI3K^H1047R^ and parental cells) of the combination. These combinations were subsequently tested in the validation experiments. **D.** Box plot comparing the measured anti-selectivity of the drug combinations predicted to anti-selective (panel B) or non-selective (panel C). Each point represents the mean anti-selectivity of one drug combination, which were each tested in 8 replicates. The difference is highly significant (Wilcoxon signed-rank test *p* < 10^−7^). **E.** Comparison of the measured anti-selectivity of IGFRi mono treatment, indicated by the horizontal gray line, with the selected IGF1Ri-containing 3-drug combinations (red box plot). IGF1Ri containing combinations are significantly more anti-selective than IGF1Ri mono treatment (one-sample t-test *p* < 10^−7^). **F.** Box plots comparing cell-viability of parental and PI3K^H1047R^ cells of the 11 (out of 17) IGF1Ri containing drug-combinations that are significantly more anti-selective than IGF1Ri mono treatment.

To do this, for all possible 3-drug combinations we optimized the concentrations such that the viability of the PI3K^H1047R^ mutants is minimized, under the constraint that the viability of the parental cells remains above 0.8 relative to DMSO control (c.f. Materials and Methods, Equation 7). To look for low-dose drug combinations, we added the constraint that each drug can be used maximally at its IC_10_. However, no drug-combination was predicted to be selective for the PI3K^H1047R^ cells at any combination of concentrations. Since none of the single drugs shows selectivity towards the PI3K^H1047R^ cells (Figure 2C), this is not very surprising. Moreover, our network reconstructions indicated that the main effect of the PI3K^H1047R^ mutation is to render the MCF10A parental line independent of growth-factor stimulation. Indeed, when we grew MCF10A parental and PI3K^H1047R^ cells in the media without growth-factor, this is what we observed (Figure S5).

To nonetheless validate our computational approach, we then looked for drug-combinations that we predicted to be anti-selective, i.e. be more effective in the parental cells than in the PI3K-mutants. In our optimizations, we found 30 such combinations with an anti-selectivity > 0.1 (Figure 4B). Interestingly, IGF1R inhibition was part of all of the 17 combinations that we predict to be most anti-selective, while its anti-selectivity in the training data was only modest (Figure 2C). However, the difference in signaling response, and specifically pAKT, was much more pronounced (Figure 2B), and this latter aspect gets picked up in the network reconstructions (Figure 3A). A particularly interesting example is the combination IGF1Ri + PI3Ki + GSK3i. Here, both PI3Ki and GSK3i at their lower dose (IC_50_) show no anti-selectivity, yet this combination is predicted to be one of the more anti-selective ones. (Figure 5B, highlighted). As a control, we also selected 44 combinations that we predicted to be non-selective for either cell line (Figure 5C). A conservative power analysis, based on the accuracy of the viability predictions and the effect size of the anti-selectivity predictions, indicated a power of 90% to detect an overall difference in selectivity between the anti-selective and control combinations.

We then treated the parental and PI3K^H1047R^ cells with the 30 predicted to be anti-selective and 44 control combinations and measured their viability (Table S5). Combinations that we predicted to be anti-selective were indeed so, and this was highly significant when compared to the non-selective control combinations (Wilcoxon signed-rank test *p* < 10^−7^, Figure 5D). Individually, 25 of the 30 combinations predicted to be anti-selective were indeed significantly so (one-sided t-test *p* < 0.05, Table S6).

As mentioned above, of the 30 predicted-to-be-anti-selective combinations we tested, 17 contain the IGF1R inhibitor, which is also mildly anti-selective as monotherapy. (None of the other inhibitors showed anti-selectivity as a monotherapy at their IC_10_, Figure S6). To rule out the possibility that our result is mainly driven by the anti-selectivity of IGF1Ri mono-therapy, we compared the 17 IGF1Ri containing drug combinations with IGF1Ri monotherapy. Figure 5E shows that each of the IGF1Ri containing combinations we tested (red box plot) is more anti-selective than IGF1Ri treatment alone (indicated by horizontal gray line). This effect is highly significant (one-sample t-test *p* < 10^−7^). When looking at the individual drug combinations, we found that 11 of the 17 IGF1Ri containing combination treatments are significantly more anti-selective than IGF1Ri monotherapy (one sided t-test *p* < 0.05, Figure 5E, Table S7). This also includes the IGF1Ri + PI3Ki + GSK3i combination highlighted above, which is the second most anti-selective combination when ranked by effect size.

These results indicate that our pipeline is capable of making an accurate prioritization of mutation-specific low-dose multi-drug combinations. Importantly, these predictions are not always obvious, and would not have been possible without the help of mathematical models of the signal transduction networks and their relation to cell viability.

## Discussion

In this study, we have shown that it is possible to predict which multiple low dose (MLD) 3-drug combinations are likely to have a mutant specific impact based on a combination of single and 2-drug high dose drug response measurements and mathematical modeling. We have used drug-perturbation experiments to reconstruct, quantify and compare signal-transduction networks of an isogenic cell line pair, and linked the responses of these networks to cell viability. No single signaling readout alone is highly predictive, but a non-linear model combining the response of ERK and AKT gave a good fit. Importantly, this showed that the short-term signaling response is predictive for cell viability, which is measured in longer-term experiments. Based on the so-obtained models we were able to predict and validate drug combinations that are were specifically effective in one cell line but not another, even though the differences between the cell lines are subtle.

One of our aims was to identify selective drug combinations, i.e. combinations that inhibit cells with an oncogenic PI3K^H1047R^ mutation more strongly then to their parental counterparts. However, according to our model, no such drug combination exist for this particular model system. The absence of oncogene-specific sensitivities is presumably due to an absence of “oncogene addiction” [1] to the PI3K mutation (or any other) in the PI3K^H1047R^ MCF10A cells. In the absence of drug-treatment the mutation has no effect on proliferation under the growth conditions we used, and this mutation therefore presumably does not induce any vulnerabilities in this cell line. Our network reconstruction suggests that the main effect of the PI3K^H1047R^ mutation on MCF10A cells is to make them growth-factor independent, consistent with previous observations [30]. Hence, the inability to identify selective drug combinations is due to the particularities of the MCF10A isogenic cell line pair model, and not due to the computational model.

While isogenic cell line pairs with a mutation knocked in are attractive models because they allow study of the effect of the mutation in isolation, they may thus not always be the best model to study oncogene-specific sensitivities. An interesting alternative approach might be to use cancer cell lines of which one of the driver mutations is removed [31–33]. Alternatively, a larger, more heterogeneous panel of cell lines with and without a particular biomarker could be used [19, 22, 34, 35]. In this scenario, one would look at commonalities in the signaling network response of the cell lines with the biomarker compared to the lines without it, and use this to propose combinations that are selective of the biomarker carrying cell lines. Finally, the use of matched tumor and normal organoids from the same patient could be used for truly personalized models.

Our aim more generally was to develop a combined experimental and computational pipeline to prioritize drug combinations that have a biomarker specific effect, and in this we did succeed. The majority of the combinations that we predicted to be anti-selective indeed were so in validation experiments. In fact, we succeeded in validating our predictions despite the fact that the differences between the cell lines in the training data were very subtle. It is to be expected that it will be easier to find mutation specific drug combinations when the effect of a mutation on the signaling networks and on cell viability is stronger.

All of the most strongly anti-selective drug combinations we identified contained the IGF1R inhibitor, but as monotherapy low-dose IGF1Ri is only mildly anti-selective. More generally, which multi-drug combinations are most selective or anti-selective is often far from obvious. For instance, while in the training data PI3Ki and GSK3i at their lower dose (IC_50_) individually show no selectivity towards the parental cells at all, the combination IGF1Ri + PI3Ki + GSK3i is one of the most selective drug combinations, both as predicted by our model and as measured validation experiments. This underscores the need for mathematical modelling in prioritizing promising combinations.

In conclusion, here we have shown that it is feasible to make accurate, non-trivial predictions about (anti-)selectivity of multi-drug combinations based on mathematical models of signaling transduction networks. In combination with suitable model systems, this framework makes it possible to rationally design biomarker-selective low-dose multidrug combinations.

## Materials and Methods

### Cells and cell culture

Human parental and PI3K^H1047R^/+ MCF10A cell lines were obtained from Horizon discovery (HD PAR-003 and HD 101-011). Cells were cultured in DMEM/F-12 including 2.5 mM L-glutamine and 15 mM HEPES, supplemented with 5% horse serum, 10 μg/mL insulin, 0.5 μg/mL hydrocortisone and 0.1 μg/mL cholera toxin. Mycoplasma tests were performed every 2 months.

### Reagents and compounds

The following inhibitors were used in this study: EGFRi (Gefitinib), IGF1Ri (OSI-906), RAFi (LY3009120), MEKi (Trametinib), ERKi (SCH772984), GSK3i (3F8), PI3Ki (BKM120), AKTi (MK-2206), mTORi (AZD8055). All inhibitors were purchased from MedKoo Biosciences. The luminex antibodies against CREB1^S133^, EGFR^Y1068^, ERK1^T202/Y204^, GSK3^S21/S9^, MEK1^S217/S221^, p70RSK^T389^, PRAS40^T246^ and RPS6^S235^ were purchased from ProtATonce Ltd. The luminex antibody against AKT1^T473^ was purchased from BioRad.

### Drug perturbation and validation experiments

All the cell-viability measurements were performed in biological triplicates, each with 2 technical replicates, using black-walled 384-well plates (Greiner 781091). Cells were plated at the optimal seeding density (200 cells per well) and incubated for approximately 24 hours to allow attachment to the plate. Drugs were then added to the plates using the Tecan D300e digital dispenser. 10 μM phenylarsine oxide was used as positive control (0% cell viability) and DMSO was used as negative control (100% cell viability). Three days later, culture medium was removed and CellTiter-Blue (Promega G8081) was added to the plates. After 2 hours incubation, measurements were performed according to manufacturer’s instructions using the EnVision (PerkinElmer). Viabilities were normalized per cell line according to (treatment − PAO_mean_)/(DMSO_mean_ − PAO_mean_). IC_50_ and IC_90_ values were fitted using the R-package MixedIC50 [36] (code available at https://github.com/NKI-CCB/MixedIC50).

The signaling response measurements were performed using 6-well plates (Greiner 657165). 300K cells per well were plated and incubated for approximately 24 hours to allow attachment to the plate. Drugs were then added to the plates and protein was harvested after 2 hours using the Bio-Plex Pro Cell Signaling Reagent Kit (BioRad 171304006M) according to the manufacturer’s instructions. Protein concentration of the samples was normalized after performing a Bicinchoninic Acid (BCA) assay (Pierce BCA, Thermo Scientific), according to the manufacturer’s instructions. Cell lysates were analyzed using the Bio-Plex Protein Array system (Bio-Rad, Hercules, CA) according to the suppliers protocol as described previously [19]. Intensities were normalized by subtracting blanks for each epitope and correcting for protein concentration.

### Computational pipeline and data analysis

#### Comparative network reconstruction

MAPK and AKT signaling networks of the parental and PI3K^H1047R^ mutant cell lines were reconstructed based on the Luminex drug-response data using Comparative Network Reconstruction (CNR)[17]. Briefly, CNR is a network reconstruction method based on Modular Response Analysis [37]. It links the matrix of measured node responses to a set of perturbations, **R**^*x*^ (where 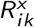 is defined as log_2_ fold change of node *i* in response to perturbation *k* in cell line *x*) to the matrix unobserved interaction strengths **r**^*x*^ (where 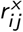 is the logarithmic partial derivative of node *i* with respect to node *j* in cell line *x*) and direct perturbation effects **s**^*x*^ (with 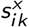 the scaled direct effect of perturbation *k* on node *i* in cell line *x*). These matrices are related through

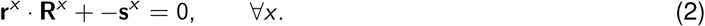

In principle, **r**^*x*^ and the values of the elements in **s**^*x*^ (the targets of the inhibitors is assumed to known) can be obtained solving this set of equations, but in practice it is often under-determined. CNR solves problem by reformulating it as optimization procedure to find a model that balances data-fit with a model complexity by penalizing the number of edges (non-zero entries in **r**) and differences between cell lines (entries in **r** that are quantitatively different between the cell lines). The optimization problem reads:

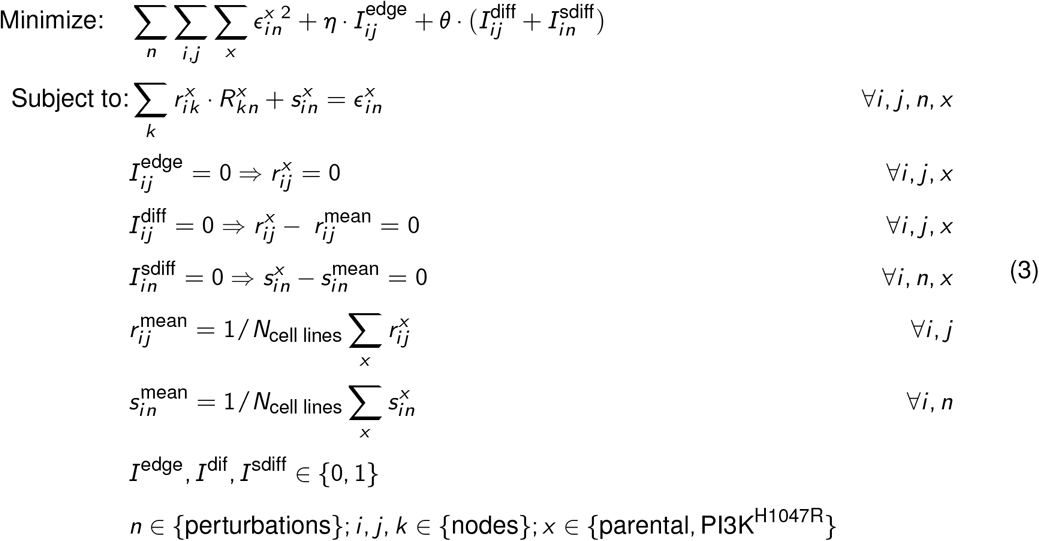

where the *ϵ*s are the model residuals. Solving this optimization problem gives the matrices **r** and **s** from a given **R**.

Additional constraints reflecting the experimental design were added to the CNR problem.

- *s_ik_* is negative and stronger for higher drug concentrations, i.e. 0 > *s_ik_*([*IC*_50_]) > *s_ik_*([*IC*_90_]).
- Each inhibitor-target pair has a single indicator for the difference in perturbations strengths for both inhibitor concentrations, i.e. if 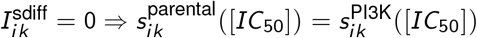 and 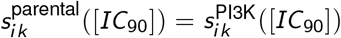.
- Most inhibitors are modelled as a perturbation to their direct target, i.e. EGFRi, MEKi, ERKi, GSK3i and AKTi are modelled as perturbations to EGFR, MEK1, ERK1, GSK3 and AKT1 respectively.
- The MEK inhibitor interferes not only with MEK phosphorylation, but also its catalytic efficiency. Hence, MEK inhibition was additionally modelled also modelled as perturbation to it’s downstream proteins (c.f. [17]).
- Some inhibitors target kinases that were not measured in our assay. The effect of these inhibitors was modelled as a perturbation to the (canonical) downstream nodes of the kinases being inhibited. Specifically, IGF1R inhibition was modelled as a perturbation to MEK1 and AKT1, PI3K inhibition as a perturbation to AKT1, RAF inhibition as a perturbation to MEK1, and mTOR inhibition as a perturbation to AKT1 and p70S6K.

Prior information about network topology was provided setting the indicators of a set of canonical MAPK and PI3K pathway interactions 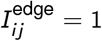. These indicator constraints are added to the optimization problem described in Equation 3. The corresponding edge-strengths, together with those of edges that might be added to the network, are found by solving the optimization problem. Hyperparameter were set to *η* = 0.1 and *θ* = 2.0 based on a leave one out cross validation loop. Single drug treatments were not included in the leave one out cross validation because each drug concentration needs to be present in at least one perturbation to estimate the corresponding parameter. The final model was obtained by restricting the topology to the prior network information with addition of the 4 edges that were identified in the leave one out cross-validation, and then performing the optimization with *θ* = 2.0.

The full Comparative Network Analysis can be found in the Jupyter notebook under the following link: https://github.com/evertbosdriesz/cnr-selective-combos/blob/master/python/cnr-mcf10a-pi3k.ipynb

#### Randomized models

To obtain the distribution of residuals for random network topologies shown in Figure 3C, 1000 models with a random topology were generated by randomly selecting 16 (out of all possible 72) edges setting the corresponding indicator to 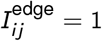 and setting the indicators of all other edges were set to 0. To focus on the effect of model topology only, we did not allow for any difference between the cell lines by setting *θ* to infinity using `cplex.infinity`. Subsequently, edge weights were obtained by solving the CNR optimization problem described in Equation 3 and corresponding RMS of residuals, defined as 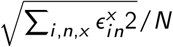, was calculated. To make a fair comparison, we also calculated the residual of our actual model without any differences between the cell lines. To this end, we set the indicators of the edges of the actual model 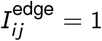 and all others to 0, set *θ* = `cplex.infinity`, and re-optimized 3.

Similarly, to obtain the distribution of residuals for random differences between the cell lines shown in Figure S2B, 1000 models with randomly selected *I*^diff^ and *I*^sdiff^ were generated. For all these models, the topology was first fixed to the topology of the actual model by setting the indicators 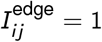 for the edges that are present in the actual model, and all others to 0. Subsequently, 13 randomly selected indicator for differences between the cell lines *I*^diff^ and *I*^sdiff^ were set to 1, the others to 0, and the optimization in Equation 7 was solved using these constraints.

#### The relation between signaling output and cell viability

The viability (relative to DMSO control) upon perturbation *k*, *v_k_*, were fitted to the following functions:

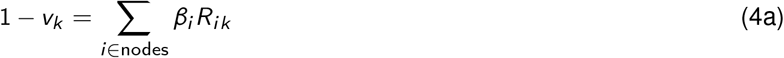

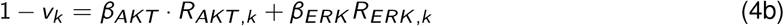

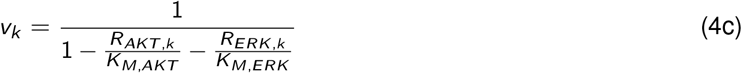

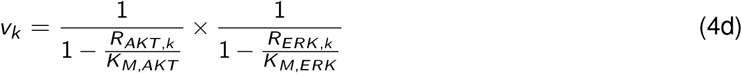

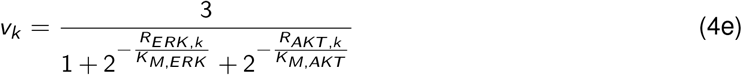

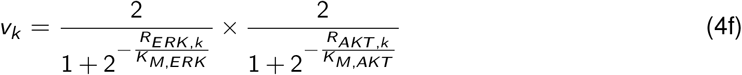

Where *R_AKT,k_* and *R_ERK,K_* are the log_2_-fold changes of pAKT and pERK relative to DMSO control upon perturbation *k*, respectively. *K_M,AKT_* and *K_M,ERK_* are the parameters to be fitted for the nonlinear equations and can be interpreted as the *R_AKT,k_* and *R_ERK,k_* values for which the viability is reduced by 50% (or 25% and 33% for equations 4e and 4f, respectively). Fitting was performed using the `lm` and `nls` functions of R [38] for the linear and non-linear models, respectively. Mean residual standard errors (*σ*) were obtained using the `sigma` function. Leave one out cross-validation was performed on a per cell-line basis. Bootstraps were performed using the function bootstrap from the ‘rsample’ package [39].

All code and details for this analysis can be found in the RMarkdown-file under the following link: https://github.com/evertbosdriesz/cnr-selective-combos/blob/master/R/02-perturbations/mapping-signaling-drugresponse.Rmd

#### Multi-drug response simulations and prediction of selective 3-drug combinations

CNR gives an estimate of the direct target inhibition of each drug only for the concentrations at which the drug was applied. To be able to simulate the effect of unseen drug concentrations, the relations between the applied concentration of drug *k*, [*I_k_*], and target inhibition of node *i* in response to this, *s_ik_* were fitted to the following function for each inhibitor-target pair,

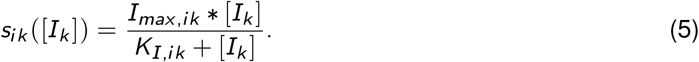

The parameters *I_max,ik_* and *K_I,ik_* were fitted to this function using the *s_ik_*-values for the [*I_k_*] = *IC*_50_ and *IC*_90_ obtained from the CNR optimizations with the `curve_fit` function from the python ‘scipy.optimize’ package [40]. For convenience all drug concentrations were normalized to the highest concentration applied (the *IC*_90_), and in all analyses only interpolations and not extrapolations are used (0 ≤ [*I*] ≤ 1).

**R**_*A+B+C*_, the vector of simulated log_2_-fold changes in response to a perturbation with 3 drugs *A, B* and *C*, at concentration [*I_A_*], [*I_B_*] and [*I_C_*] was calculated as

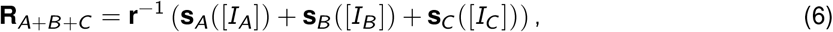

to obtain *R_AKT,A+B+C_* and *R_ERK,A+B+C_*. These were then used to calculate viability according to Equation 4. Together, this allows for simulating the effect on cell viability of drug combinations and concentrations that were not seen in the training data.

For each possible 3-drug combination, the selectivity for cell line *x* relative to *y* was optimized by solving the following optimization problem:

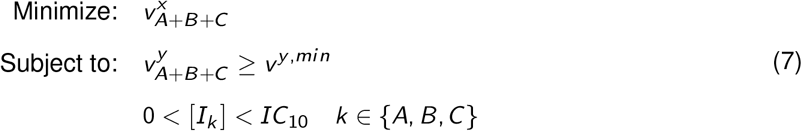

where *v^y,min^* is the cutoff used for the minimal viability that cell line *y* should have under the 3-drug treatment, and that we (somewhat arbitrarily) set to 0.8.

Similarly, unselective control combinations were obtained by solving the optimization problem:

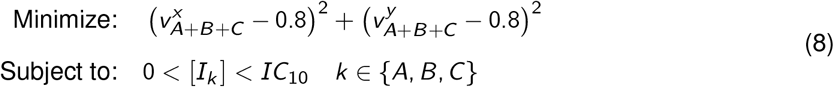

for all possible 3-drug combinations.

A power analysis of the predictions was performed by performing 1000 simulations with addition gaussian noise with a mean 0 and a standard deviation 0.25 (based on the residuals of our viability predictions) to the results, and counting in what fraction their was an observable difference between the two groups.

The optimizations were performed in Wolfram Mathematica [41] (version 12.0) using the `NMinimize` function. The full optimization and power analysis can be found in the Mathematica notebook under the following link: https://github.com/evertbosdriesz/cnr-selective-combos/blob/master/mathematica/optimize-combinations.nb

## Data and Code availability

All data and code required to reproduce the results and figures in this paper are available at https://github.com/evertbosdriesz/cnr-selective-combos.

## Acknowledgements

This work was supported by funding from the Oncode Institute and the Gravity Program CGC.nl, funded by The Netherlands Organisation for Scientific Research (NWO).

## Supplementary Figures

**Figure S1:**
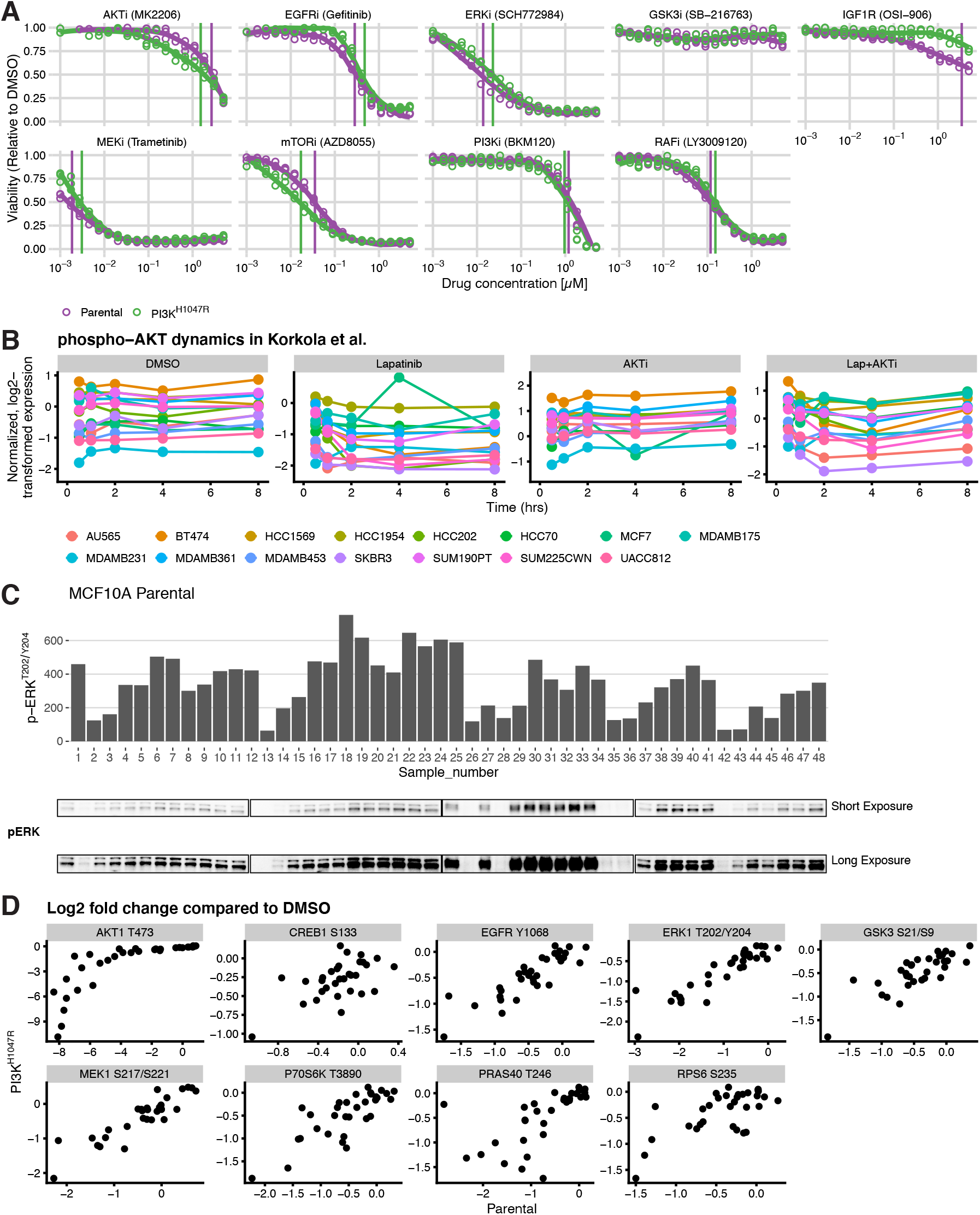
**A.** Dose-response curves of the inhibitors used in this study. **B.** Dynamics of AKT activity after PI3K pathway inhibition from Korkola *et al*. [29]**C.** Correlation between phospho-ERK quantification using Luminex (top) and Western blot (bottom). **D.** Correlation between the response in parental (x-axis) and PI3K^H1047R^ (y-axis) cells. Response is defined as log_2_-fold change compared to DMSO controls.

**Figure S2:**
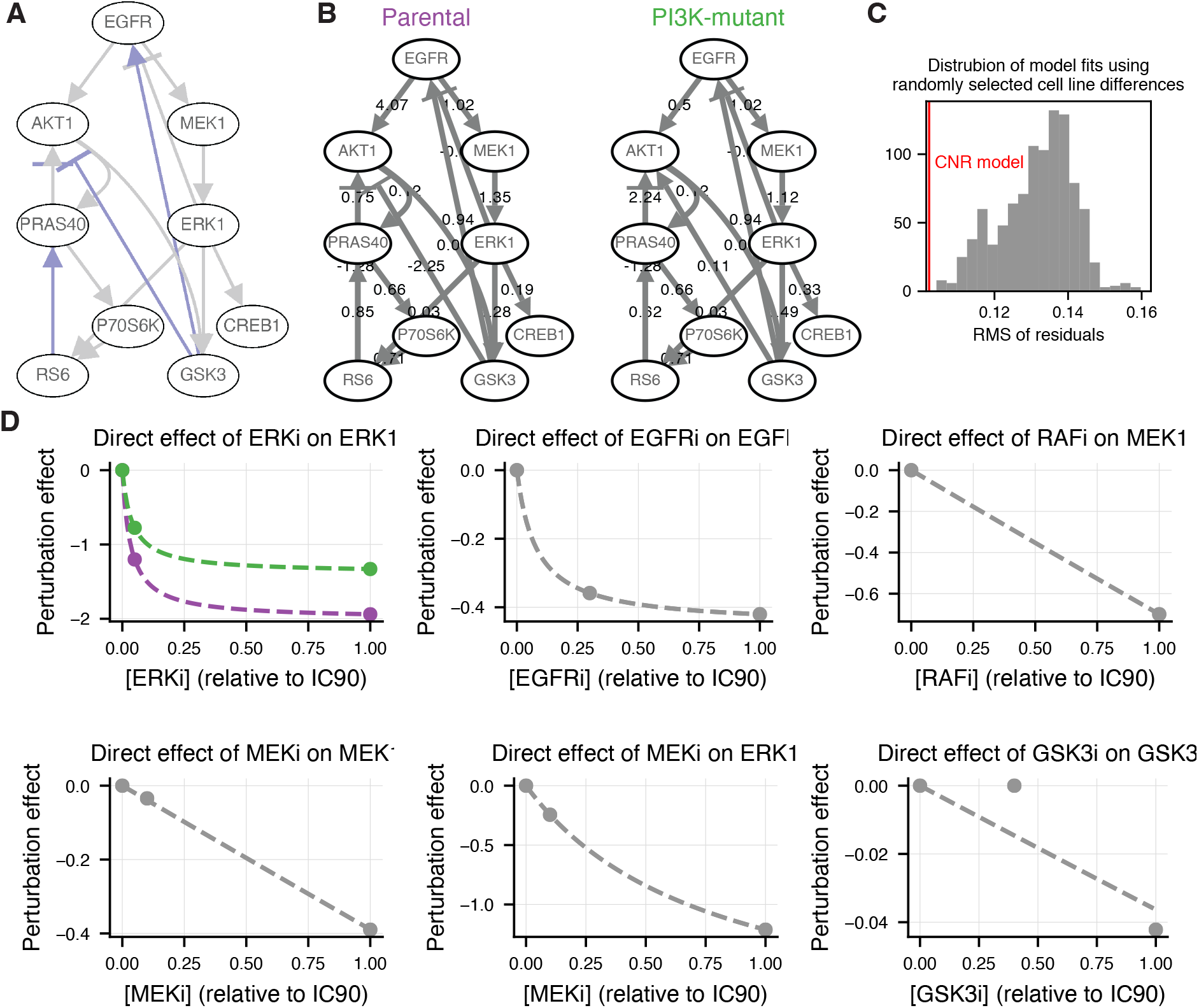
**A.** Network topology used for modeling. Edges used as prior information are indicated in gray. Edges added in a leave one out cross validation loop are indicated in purple. **B.** Network models of the parental (left) and PI3K^H1047R^ cells. Edge labels indicate reconstructed interaction strenghts (*r_ij_* terms in Equation 2 and 3). **C.** Significance of identified differences between the cell lines. Distribution of the residuals of 1000 model optimizations in which random edges were selected to allow to differ between the two cell. The selected model has a better model fit than all 1000 of these models. **D.** The estimated direct effect of different inhibitors on their target, as a function of applied inhibitor concentration (*s_ij_* terms in Equation 2 and 3). Points indicate the estimated effects obtained from the CNR reconstruction, at the concentrations used in the perturbation experiments. The dashed lines indicate the interpolated curves between these points. (c.f. Materials and Methods, Equation 5)

**Figure S3:**
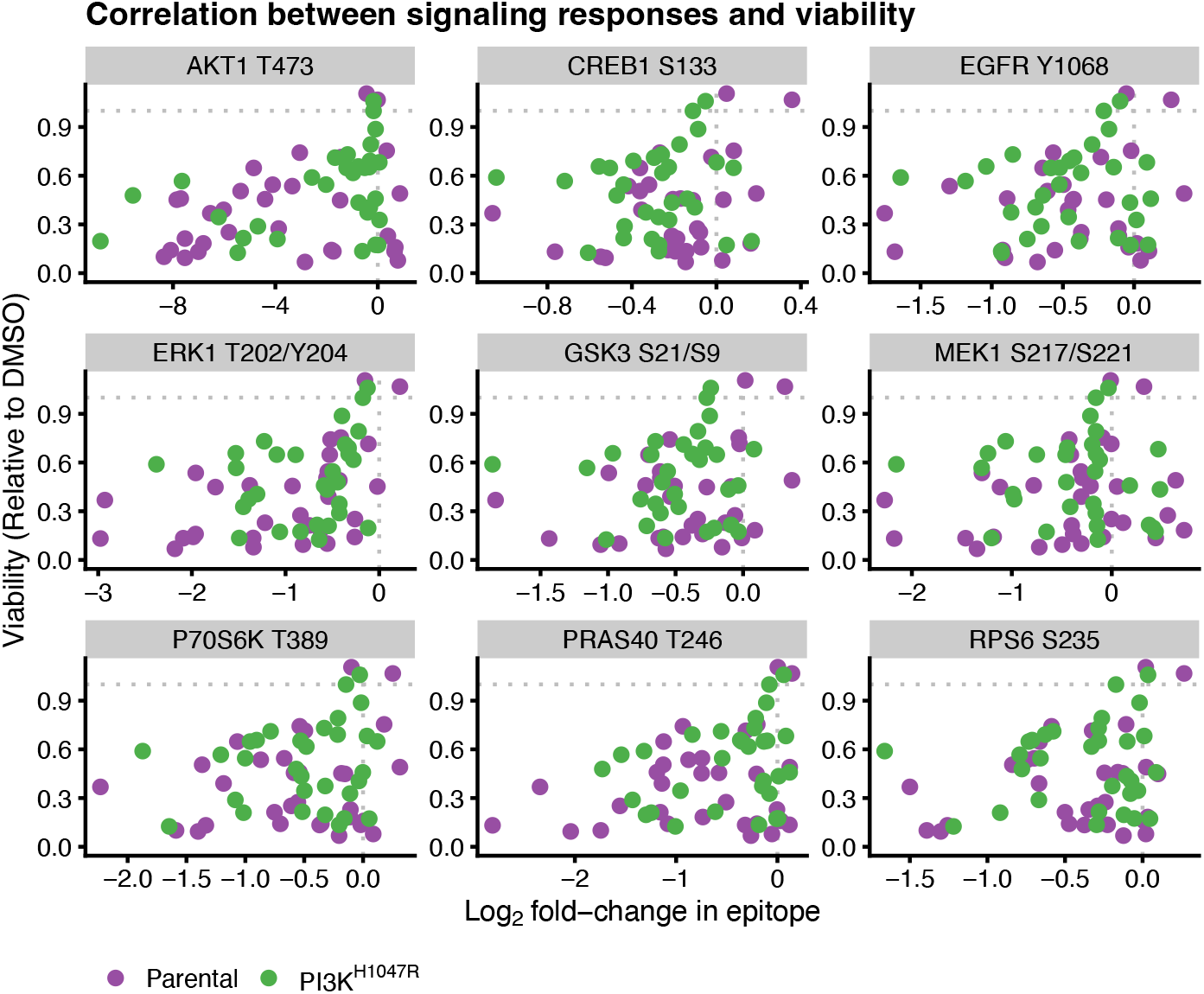
Correlation between node response and cell viability of all measured nodes.

**Figure S4:**
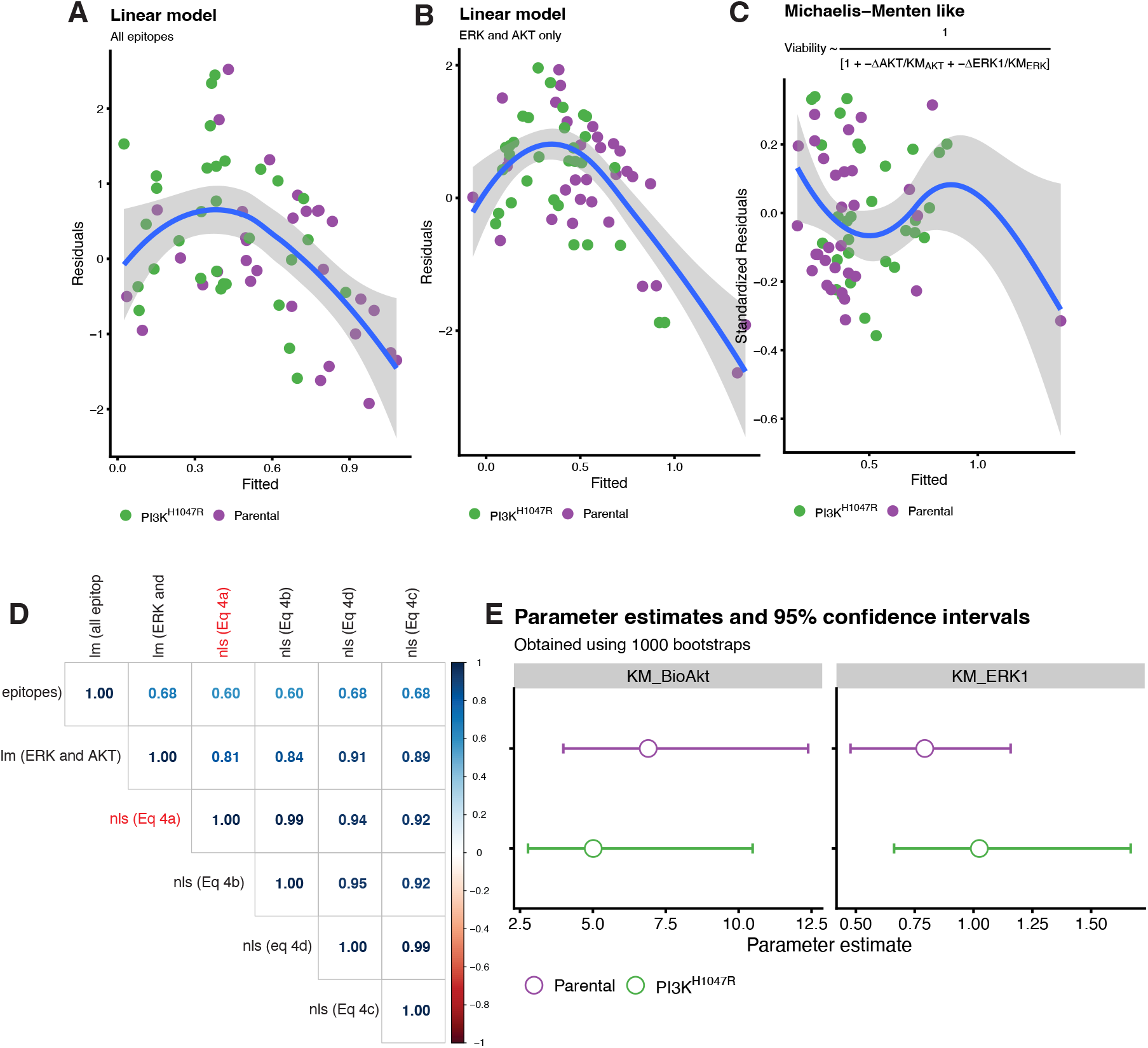
Evaluation of model fits relating signaling response to cell viability. **A-C.** Residuals as a function of fitted values for the model fits. **A.** Linear model with all epitopes as predictor. **B.** Linear model with only *R_AKT_* and *R_ERK_* as predictor. **C.** Non-linear model that gave the best fit (Equation 4c). **D.** Pearson correlation between model predictions of all models tested show that the predictions of all non-linear models are highly similar. Predictions are obtained from the leave-one-out cross-validation. The selected model (Equation 4c) is highlighted in red. The plot is generated using the corrplot-function of the corrplot R-package [42]. **E.** Bootstrapping intervals of the estimated values for the parameters K_M,AKT_ and K_M,ERK_ in Equation 4c

**Figure S5:**
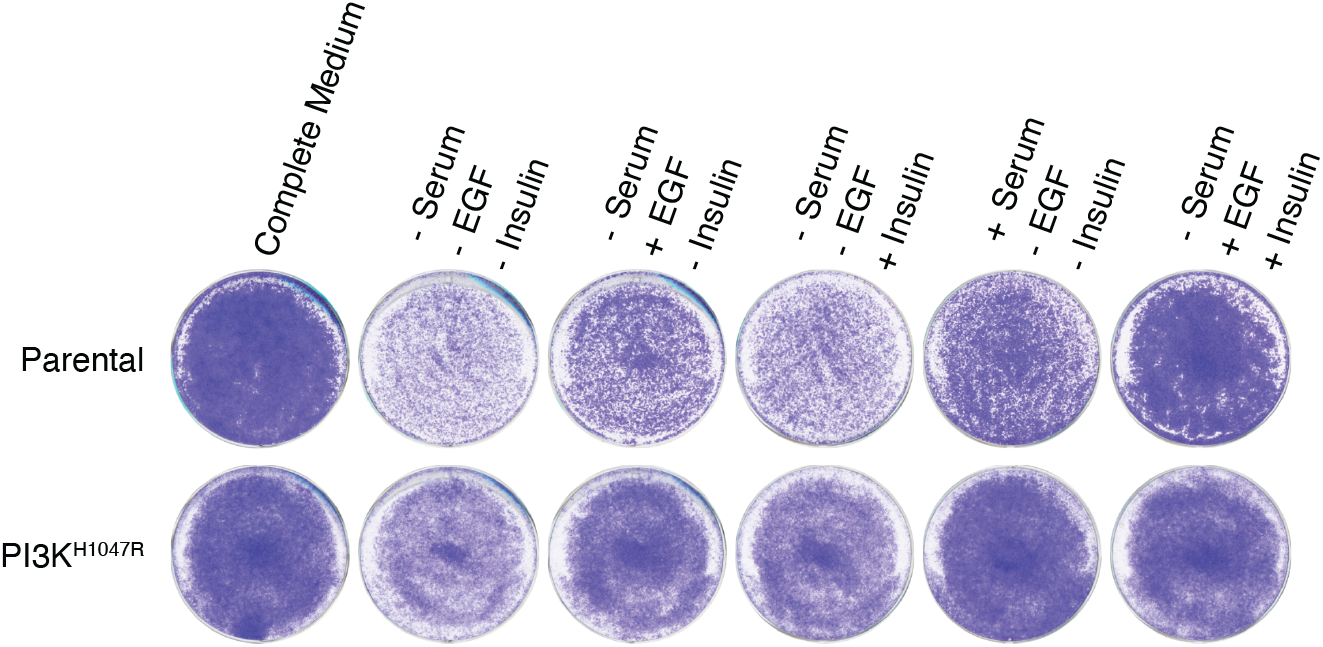
Growth of MCF10A parental and PI3K^H1047R^ cells in different growth media. In contrast to the parental cells the PI3K mutant cells grow well in the absence of serum if either Insulin or EGF is provided.

**Figure S6:**
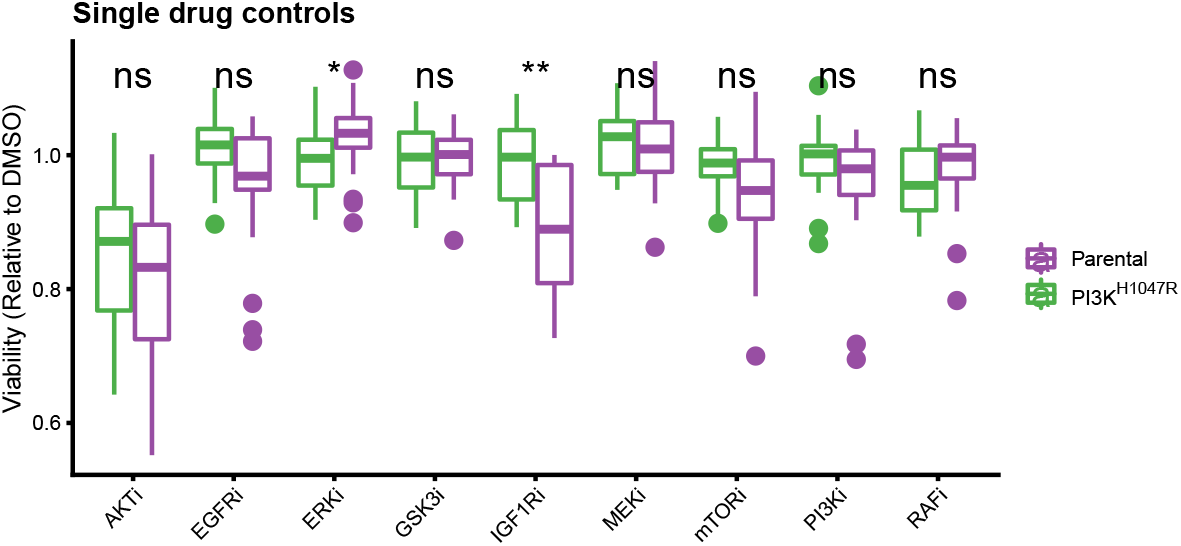
Viability of the low-dose single-drug controls, all measured at their *IC*_10_. Except for IGF1Ri, none of the drugs show selectivity towards the parental cells. Treatments were performed in 8 replicates.

